# Influence of Preservation Methods, Sample Medium and Sampling Time on eDNA Recovery in a Neotropical River

**DOI:** 10.1101/489609

**Authors:** Naiara Guimarães Sales, Owen Simon Wangensteen, Daniel Cardoso Carvalho, Stefano Mariani

## Abstract

Environmental DNA (eDNA) has rapidly emerged as a promising biodiversity monitoring technique, proving to be a sensitive and cost-effective method for species detection. Despite the increasing popularity of eDNA, several questions regarding its limitations remain to be addressed. We investigated the effect of sampling medium and time, and preservation methods, on fish detection performance based on eDNA metabarcoding of neotropical freshwater samples. Water and sediment samples were collected from 11 sites along the Jequitinhonha River, Southeastern Brazil; sediment samples were stored in ethanol, while the same amounts of water per sample (3L) were stored in a cool box with ice, as well as by adding the cationic surfactant Benzalkonium chloride (BAC). Sediment and water samples yielded a similar amount of fish MOTUs (237 vs 239 in the first sampling event, and 153 vs 142 in the second sampling event). Water stored in ice provided better results than those preserved in BAC (239 and 142 vs 194 and 71 MOTUs). While documenting the effectiveness of eDNA surveys as practical tools for fish biodiversity monitoring in poorly accessible areas, we showed that keeping water samples cooled results in greater eDNA recovery and taxon detection than by adding cationic surfactants as sample preservatives. Furthermore, by comparing two sets of samples collected from the same locations at a three-week interval, we highlight the importance of conducting multiple sampling events when attempting to recover a realistic picture of fish assemblages in lotic systems.

## INTRODUCTION

Environmental DNA metabarcoding has been hailed as a promising tool for biodiversity assessment and monitoring worldwide, in both marine and freshwater ecosystems (Bohmann et al., 2014; Boussarie et al., 2018; Deiner et al., 2017; Hänfling et al., 2016; Pont et al., 2018; Thomsen & Willerslev, 2015). This method relies on obtaining the DNA shed by organisms in the surrounding environment (e.g. water, soil), amplifying it with primers targeting the taxonomic spectrum of interest, and high-throughput sequencing it to reconstruct community composition (Bohmann et al., 2014; Handley et al., 2018; Valdez-Moreno et al., 2018; Valentini et al., 2016).

Despite the increased number of publications in the past decade, the application of eDNA techniques is still not considered straightforward (Taberlet, Bonin, Zinger, & Coissac, 2018). Molecular and bioinformatics protocols continue to be revised and optimized, while uncertainties remain as to how to streamline and rationalize sampling and sample preservation (Dickie et al., 2018). The usefulness of eDNA approaches depend on their ability to provide effective and accurate detection of species, thus requiring a better understanding of the factors influencing detection rates (Lodge, 2012). Detectability of eDNA in environmental samples is limited mainly by three processes: i) eDNA production (i.e. rate of DNA shedding), ii) degradation, iii) removal and transport (Barnes and Turner, 2016; Strickler, Fremier & Goldberg, 2015). Several factors can affect eDNA production, such as the type of organism/species (with some species showing a higher eDNA release rate than others Maruyama, Nakamura, Yamanaka, Kondoh, & Minamoto, 2014; Sassoubre, Yamahara, Gardner, Block, & Boehm, 2016), biomass, density and life stage of specimens (Maruyama et al., 2014; Takahara, Minamoto, Yamanaka, Doi, & Kawabata, 2012), season (Buxton, Groombridge, Zakaria, & Griffiths, 2017), and water oxygen and temperature which can cause behavioral and physiological changes (e.g. stress) and affect metabolic rates, hence influencing eDNA production (Maruyama et al., 2014; Pilliod, Goldberg, Arkle, & Waits, 2014). After eDNA is released in the water it starts to be removed through transport and/or degradation. eDNA molecules can settle and bind to sediment, and/or be transported by long distances depending on the type of environment (e.g. lotic, lentic), and thus, degrade and become diluted during the transport downstream (Strickler et al., 2015).

The DNA released in the environment can be degraded at a fast pace, hampering the identification of rare species and providing false negatives (Barnes et al., 2014; Dejean et al., 2011; Pilliod et al., 2014; Strickler et al., 2015), which leads to the need for improved preservation systems that can maximize eDNA recovery (Fonseca, 2018; Hansen, Bekkevold, Clausen, & Nielsen, 2018). The persistence of DNA in environmental samples can be influenced by many factors (e.g. temperature, microbial activity, pH, salinity, solar radiation), and detectability of eDNA in water has been shown to be associated with cold temperatures, alkaline conditions, and low UV-B levels (Strickler et al., 2015; Tsuji, Ushio, Sakurai, Minamoto, & Yamanaka, 2017), even though several studies suggest a negligible role of temperature, UV levels or seasonality on DNA degradation (Andruszkiewicz. Sassoubre, & Boehm, 2017; Collins et al., 2018; Robson et al., 2016).

The most recommended approach to reduce degradation is to extract the DNA as quickly as possible after sampling. However, due to the constraints of field work conducted in remote sites located far from laboratory facilities (e.g. difficulties for on-site filtration due to lack of equipment, and risk of contamination), the filtering process and subsequent DNA extraction might not be possible or advisable, and a preservation method must be employed in order to block biological activities and minimize DNA degradation.

Different approaches have been tested to preserve water samples before the filtering process, showing distinct benefits and drawbacks. Storing the samples at low temperatures, including freezing the samples or cooling using a cool box, are widely employed; however, these approaches entail equipment requirement increase; whereas the efficiency of cooling the samples has also been questioned (Eichmiller, Best, & Sorensen, 2016; Pilliod et al., 2014). Inclusion of buffers, such as EtOH–NaAc (ethanol-sodium acetate) solution, have been reported to show an eDNA persistence rate similar to samples stored in ice (Ladell, Walleser, McCalla, Erickson, & Amberg, 2018), however, when sampling larger volumes of water the increased final volume obtained (i.e. addition of over 2x of solution) might be considered as a problem during long sampling campaigns. Recently, Yamanaka et al. (2017) tested the addition of cationic surfactants as preservatives to suppress DNA degradation at ambient temperatures and demonstrated the efficiency of Benzalkonium chloride (0.01%) in retaining eDNA concentration even after 10-day incubation at 21ºC. Still, despite being considered as an effective eDNA preservative, this preservation method was restricted to a species specific eDNA recovery test and the effectiveness of the cationic surfactant in preserving eDNA samples for metabarcoding analysis has not yet been evaluated.

The application of eDNA as a biodiversity assessment tool requires the development, field validation and optimization of protocols in order to minimize bias and tailor procedures to the variety of environments and habitats investigated (Taberlet et al., 2018). Furthermore, the occurrence of a time lag between species presence and sampling event can contribute to DNA degradation leading to an erroneous inference of species absence (i.e. short time frame detection due to high degradation rates may hamper the eDNA efficiency in detecting species where they are present). Sediment samples have shown to contribute to tackling this issue once DNA attached to sediments can be detected longer than in the water column. In addition, sediment samples can provide a higher concentration and longer persistence of genetic material for studying past and current species presence, also contributing to understand issues associated with eDNA transport and removal (Turner, Uy, & Everhart, 2015).

Neotropical freshwaters harbor high, and often understudied (Sales, Mariani, Salvador, Pessali, & Carvalho, 2018), biodiversity and eDNA could assist biodiversity assessment and monitoring programs, with the ultimate aim to contribute to conservation and management strategies. Higher temperatures and solar radiation associated with increased turbidity in tropical waters might contribute to make rivers in the tropics a challenge for eDNA studies due to possibly higher degradation rates (Barnes et al., 2014; Matheson, Gurney, Esau, & Lehto, 2014; Pilliod et al., 2014). A rapid removal of eDNA (through transport and degradation) might hamper the detection of species and lead to false negatives (Hansen et al., 2018), compromising the use of this method for biodiversity assessment and monitoring. In this context, testing effectiveness of sampling methods is particularly important in remote and tropical locations (Ladell et al., 2018). Furthermore, the knowledge regarding the use of eDNA in tropical rivers remains scarce and despite being considered as a promising tool for fish biodiversity assessment in this region, this approach still requires the optimization of field and laboratory protocols (Cilleros et al., 2018). To our knowledge no study has been conducted in Neotropical catchments to evaluate the effect of sampling medium and preservation methods in lotic environments. Here we obtained water and sediment samples from 11 sites located along the main stem of River Jequitinhonha (South-Eastern Brazil), and: a) compared two preservation methods for water samples (cooling the samples using ice and adding the cationic surfactant Benzalkonium chloride – BAC); b) compared MOTU recovery from water vs sediment samples, and c) examined the influence of short-term temporal sample replication by sampling the same locations over a three-week interval.

## MATERIAL AND METHODS

### Study Site

The Jequitinhonha River Basin, located in Southeast Brazil, flows through two biodiversity hotspots (Atlantic Forest and Cerrado) encompassing an area of 70,315 km^2^ and running over 1082 km. This region is characterized by tropical climate and environmental heterogeneity, including semi-arid regions with high temperatures (annual mean of 24.9°C) and dry period extending over six months per year (Climate-Data, 2018, Bilibio, Hensel, & Selbach, 2011). This catchment, located in one of the poorest and least studied regions of Brazil, is part of an ecoregion (Coastal Drainages of Eastern Brazil) that harbors considerable fish biodiversity and one of the highest numbers of endemic and threatened fish species in Brazil (Machado, Drummond, & Paglia, 2008, Pugedo, Andrade-Neto, Pessali, Birindelli, & Carvalho, 2016, Rosa & Lima, 2008).

### eDNA sampling and processing

Sediment and water samples were obtained from 11 sample sites, in the Jequitinhonha River Basin, during two replicated sampling events carried out in January-March 2017 (Figure 1, Table S1 Supporting information). In each sampling event, 6 liters of water were collected from each sample site (i.e. 3 subsamples of 1 liter each, per treatment) and before the filtering process the water was preserved using two different methods to compare their efficiency. Upon collection, one set of samples was stored at low temperatures (using a cooling box with ice), while in the other batch the cationic surfactant benzalkonium chloride (BAC) was added at a final concentration of 0.01% (Yamanaka et al. 2017). Water samples were filtered approximately 8 hours after collection, using Microfil V, 100mL, mixed cellulose esters (MCE) filters (diameter: 47 mm, pore size: 0.45 μm, Merck Millipore) (Bakker et al. 2017; Deiner et al., 2018) in combination with an automatic vacuum pump. Filters were stored in microcentrifuge tubes containing silica beads (Bakker et al. 2017). Sediment samples (2 samples/locality) were obtained in the shores, from the superficial layer (approximately 5cm), and were stored in 50mL centrifuge tubes and preserved in 100% ethanol.

**Figure 1:**
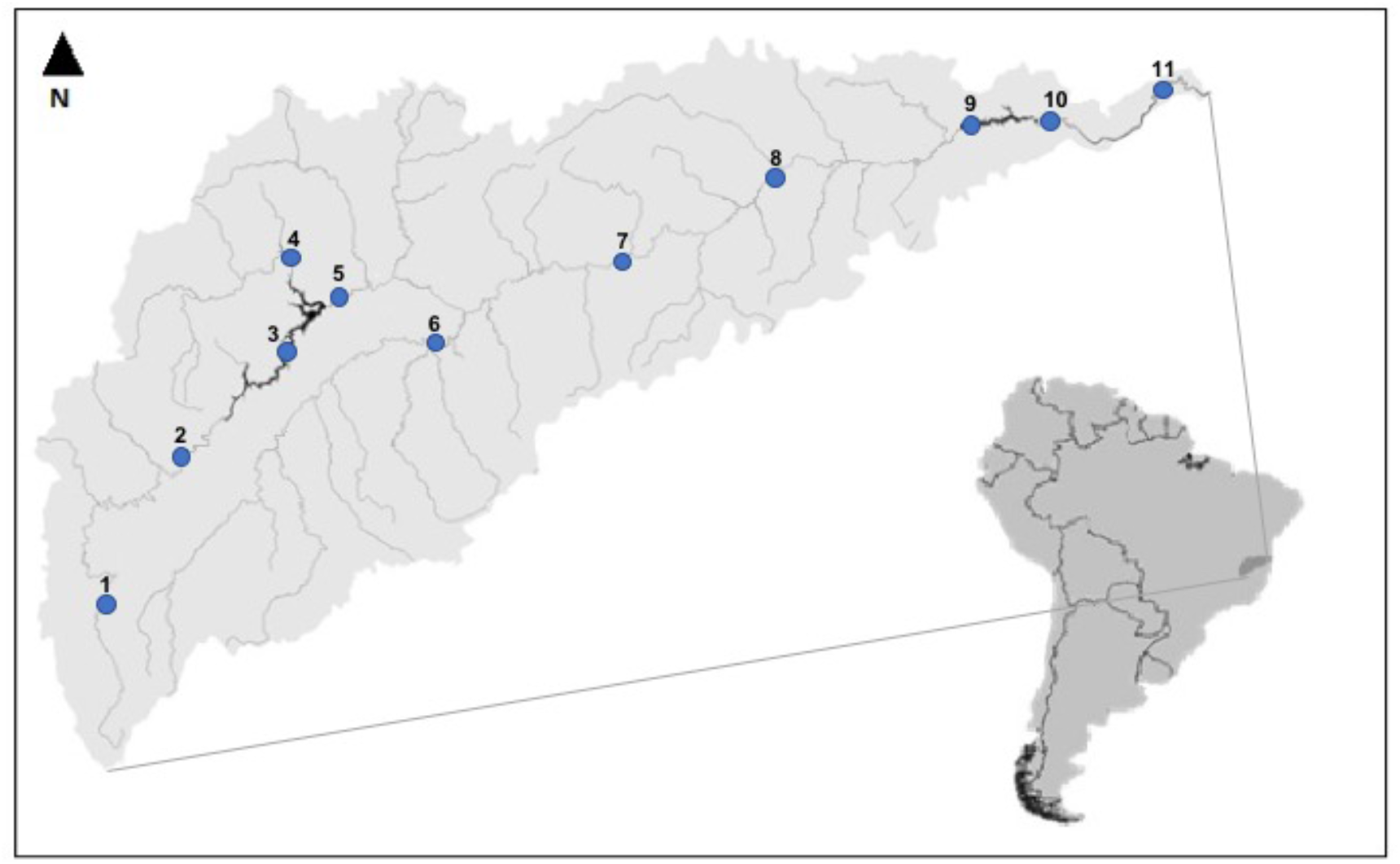
Map of Jequitinhonha river basin sampling locations.

DNA extraction from the filters was conducted using the DNeasy PowerWater Kit (Qiagen) and DNA from the sediments was extracted from 10g of sediment using DNeasy PowerMax Soil Kit (Qiagen), following the manufacturer’s protocol. Purified extracts were checked for DNA concentration in a Qubit fluorometer (Invitrogen).

A contamination control procedure was applied in both field and laboratory works to avoid the occurrence of contamination. All samples were stored in sterile collection bottles, disposable gloves were worn at all times, sampling and laboratory equipment and surfaces were treated with 50% bleach solution for 10 minutes, followed by rinsing in water after each use. Filtration blanks were run between every sample site, immediately before the next filtration in order to test for potential contamination during the filtration stage.

### Amplification, Library preparation and sequencing

The amplification of eDNA metabarcoding markers was conducted using a previously published fish-specific 12S primer set (Miya et al., 2015). Amplicons of ∼172bp from a variable region of the mitochondrial 12S rRNA gene were obtained with the primers (MiFish-U-F, 5′-GCCGGTAAAACTCGTGCCAGC-3′; MiFish-U-R, 5′-CATAGTGGGGTATCTAATCCCAGTTTG-3′).

A total of 183 samples including collection blanks and laboratory negative controls were sequenced in a single multiplexed Illumina MiSeq run using 2 sets of 96 primers with seven-base sample-specific oligo-tags and a variable number (2-4) of leading Ns (fully degenerate positions) to increase variability in amplicon sequences. PCR amplification was conducted using a single-step protocol and to minimize stochasticity in individual reactions, PCRs were replicated three times for each sample and the products subsequently pooled into single samples. The PCR reaction consisted of a total volume of 20 µL including 10 µl Amplitaq; 0.16 µl of bovine serum albumin; 1 µl of each of the two primers (5 µM); 5.84 µl of ultra-pure water and 2 µl of DNA template. The PCR profile included an initial denaturing step of 95°C for 10 min, 40 cycles of 95°C for 30s, 60°C for 45s, and 72°C for 30s and a final extension step of 72°C for 5 min. Amplifications were checked through electrophoresis in a 1.5% agarose gel stained with GelRed (Cambridge Bioscience). PCR products were pooled in two different sets and purified using MinElute columns (Qiagen), and Illumina libraries were built from each set, using a NextFlex PCR-free library preparation kit (Bioo Scientific) with unique 6-bp library tags. A left-sided size selection was performed using 1.1x Agencourt AMPure XP (Beckman Coulter). Libraries were then quantified by qPCR using a NEBNext qPCR quantification kit (New England Biolabs) and pooled in equimolar concentrations along with 1% PhiX (v3, Illumina). The libraries were run at a final molarity of 10pM on an Illumina MiSeq platform in a single MiSeq flow cell using the 2x 150bp v2 chemistry.

### Bioinformatics analyses

Bioinformatic analyses were based on the OBITools metabarcoding package (Boyer et al. 2016). FastQC was used to assess the quality of the reads, paired-end reads were aligned using illuminapairedend, and dataset demultiplexing and primer removal were then conducted using ngsfilter command. A bespoke filter using obigrep was used to select fragments of 140–190bp and remove short fragments originated from library preparation artefacts (primer-dimer, non-especific amplifications) and reads containing ambiguous bases. Clustering of strictly identical sequences was performed using obiuniq and a chimera removal step was applied in vsearch (Rognes, Flouri, Nichols, Quince, & Mahé, 2016) through the uchime-denovo algorithm (Edgar, Haas, Clemente, Quince, & Knight, 2011). Molecular Operational Taxonomic Unit (MOTU) delimitation was performed using SWARM 2.0 algorithm (Mahé, Rognes, Quince, de Vargas, & Duthorn, 2015) with a distance value of d=3 (Siegenthaler et al., 2018) and ecotag (Boyer et al. 2016) was used for the subsequent taxonomic assignment, with a custom reference database including all known vertebrate sequences for the sequenced 12S fragment (Siegenthaler et al., 2018). Ambiguous taxonomic assignments after ecotag were checked using BLAST against the Genbank nucleotide database.

A conservative approach was applied to our analyses to avoid false positives and exclude MOTUs/reads putatively belonging to sequencing errors or contamination. Reads detected in the negative controls were removed from all samples, and MOTUs containing less than 5 reads were excluded from subsequent analyses.

### Statistical analyses

Samples were grouped according to the treatments analyzed (Table 1) and afterwards all statistical analyses were performed in R v3.5.1 (https://www.R-project.org/). Due to differences in the sequencing depth for each sample, relative read abundances were used for all statistical analyses (i.e. for each sample the MOTU counts were divided by the total amount of reads). The vegan package was used to perform the nonparametric method Permutational multivariate analysis of variance (PERMANOVA) (Anderson, 2017), through the ‘adonis’ function (Bray-Curtis dissimilarities, 1000 permutations). Comparisons were performed on relative abundances calculated for MOTUs in each sample site, per preservation method (BAC vs ICE), sampling time (1st round vs 2nd round), and per sampling medium (water vs sediment), to verify the influence of these factors over eDNA recovery. A significance threshold of p < 0.05 was applied at all analyses.

**Table 1:** Treatments analyzed according to sampling medium, preservation method used and sampling event.

| CODE | Sampling Medium | Preservation method | Sampling event |
| --- | --- | --- | --- |
| SED1 | Sediment | Ethanol | 1 |
| SED2 | Sediment | Ethanol | 2 |
| BAC1 | Water | Benzalkonium chloride | 1 |
| BAC2 | Water | Benzalkonium chloride | 2 |
| ICE1 | Water | ICE | 1 |
| ICE2 | Water | ICE | 2 |

Non-metric multidimensional scaling plots were obtained using Bray-Curtis dissimilarity, through PAST3 software (Hammer, Harper, & Ryan, 2001). ggplot2 and esquisse packages were used to build ggplot charts in R, and due to an incomplete reference database and a relatively low taxonomic resolution of the 12S fragment we used the taxonomic assignment down to family level to compare those methods regarding their performance in detecting teleost fish communities. Venn diagrams were obtained with BioVenn (Hulsen, Vlieg, & Alkema, 2008).

## RESULTS

### Library quality and raw data

A total of 16,104,492 raw reads were obtained in one Illumina MiSeq run (Library 1: 6,399,823 reads, Library 2: 9,704,669 reads), including 44 sediment samples and 132 water samples. 10,064,034 reads were kept after initial quality filtering and removal of chimaeras. After applying a subsequent conservative filtering step (retaining only reads taxonomically assigned to Actinopterygii, and removal of MOTUs containing less than 5 reads) the number of reads per sample ranged from 0 (sample 10 – sediment; second sampling event) to 127,250. The final dataset comprised 311 MOTUs distributed differently in each treatment analyzed (Figure 2, Table 2).

**Figure 2:**
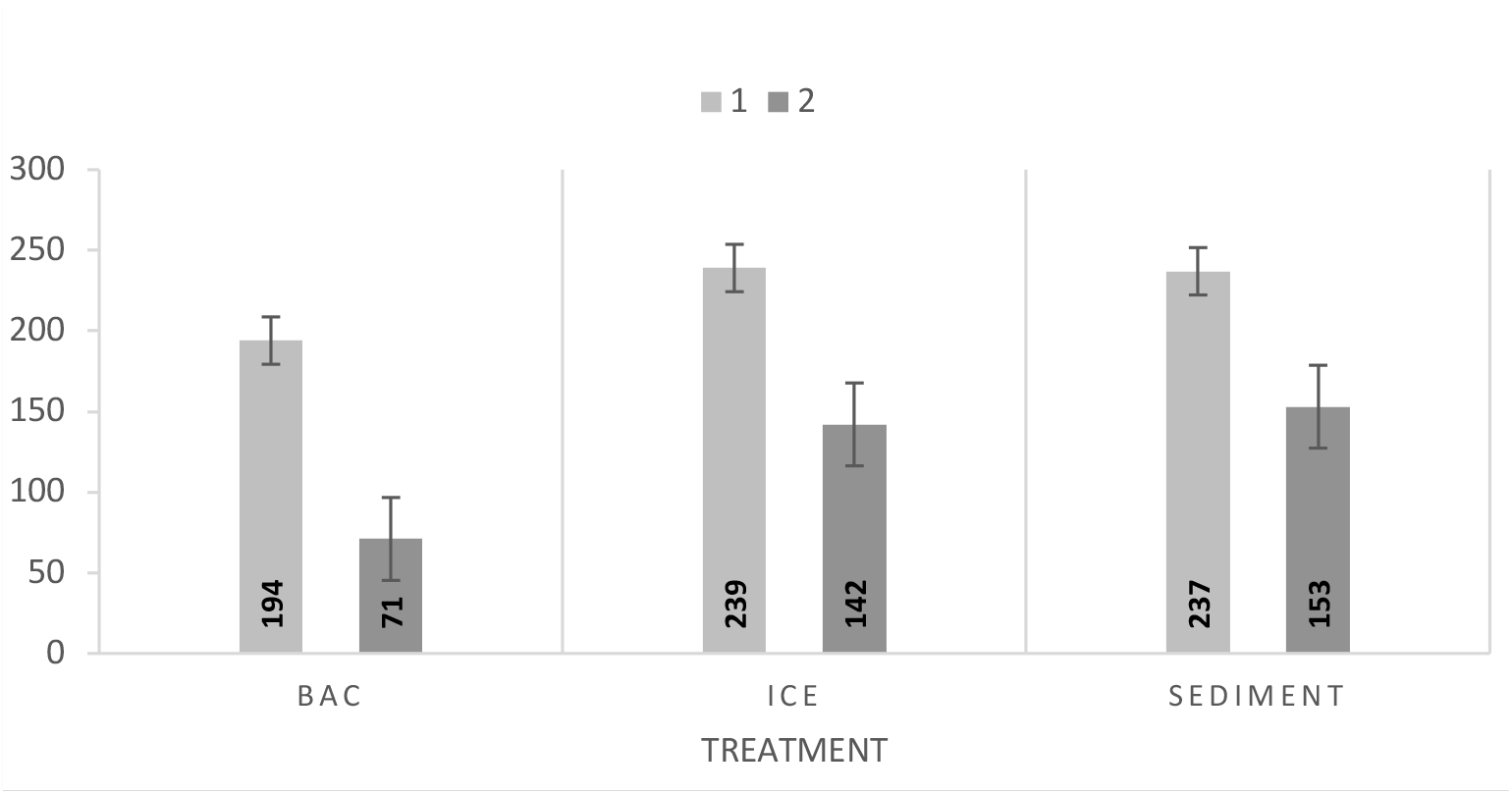
Total number of MOTUs recovered per sampling medium and preservation method (sediment vs water – BAC and ICE) and sampling event.

**Table 2:** MOTUs recovery per sampling medium, preservation method, and sampling event.

| Sampling medium | Preservation method | Sampling event | MOTUs |
| --- | --- | --- | --- |
| Water | BAC | 1 | 194 |
|  |  | 2 | 71 |
|  | ICE | 1 | 239 |
|  |  | 2 | 142 |
| Sediment | ETHANOL | 1 | 237 |
|  |  | 2 | 153 |

### Taxonomic assignment

All MOTUs from the sediment samples could be taxonomically assigned at order level (see Appendix S1, Supporting information) whereas at family level the assignment rate was 96.4% (SED1) and 95.68% (SED2). Regarding the water samples, at order and family levels the assignment rates were, respectively, 98.97% and 95.88% for BAC1, 97.47% and 93.68% for BAC2, 100% and 96.83% for ICE1, and 98.72% and 94.17% for ICE2.

### Influence of preservation method, sampling medium, and sampling time

All results of the PERMANOVA analyses (Bray-Curtis, p<0.005), including effect size (R^2^) and significance (p-value) are summarized in Table S2, Supporting information. A significant difference (p<0.05) in MOTU composition among all the treatments was found and to verify the influence of preservation methods, sampling medium, and sampling time we performed pairwise comparisons for all combinations of treatments.

The influence of preservation method on MOTU diversity recovery was small (around 2% variance explained) but significant between samples collected during the first sampling event (BAC1 vs ICE1, p=0.016). However, no significant effect was detected for the preservation methods in the second sampling event (BAC2 vs ICE2, p=0.06) (Table S2).

Overall and also in all pairwise comparisons, a significant difference between sediment and water samples was detected. Non-metric multidimensional scaling (nMDS) (Figure 3) showed a much greater variability among the water samples when compared to the sediment ones, and a greater separation of samples was apparent for the first sampling event (Figure 3A). During the second sampling, a higher similarity between sediment and water samples preserved cooled was found (Figure 3B), and the highest effect size (R^2^=0.08) was found between SED2 and BAC2 (sediment and water samples preserved in BAC, collected during the second sampling event).

**Figure 3:**
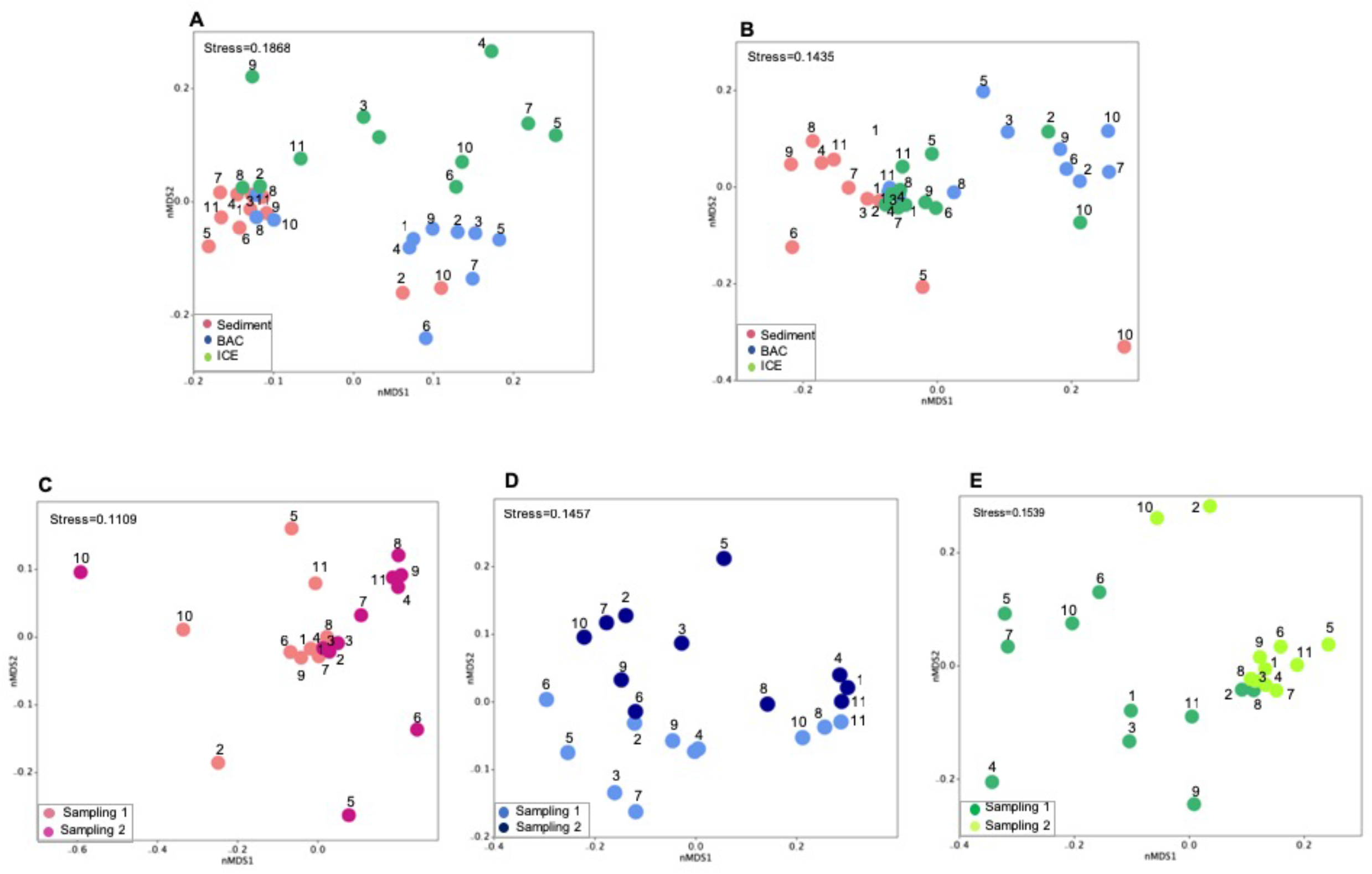
Non-metric multidimensional scaling (nMDS) plots showing similarities of sample sites per sampling event. Analyses based on A) Sampling event 1; B) Sampling event 2; C) Sediment samples; D) Water samples preserved using BAC; and E) Water samples preserved using ICE.

When testing for the effect of sampling event, the community composition differed from the two events for all treatments analyzed, showing a highest effect size for the sediment samples (R^2^=0.07) and a lower effect size for the water samples preserved in BAC (R^2^=0.04). A smaller effect was found for preservation method than sampling medium and time. Despite showing significant differences, overall, the R^2^ effect sizes never accounted for any more than 8% of the variance, with a mean around 6%.

The Venn diagram overlaps showed a high similarity between the treatments in the first sampling event with 56.78% of the MOTUs detected in all of them (Figure 4). However, for the second sampling event a higher dissimilarity was detected when comparing the methods applied with only 27.55% of the MOTUs recovered being detected in all three methods (sediment, BAC, ICE).

**Figure 4:**
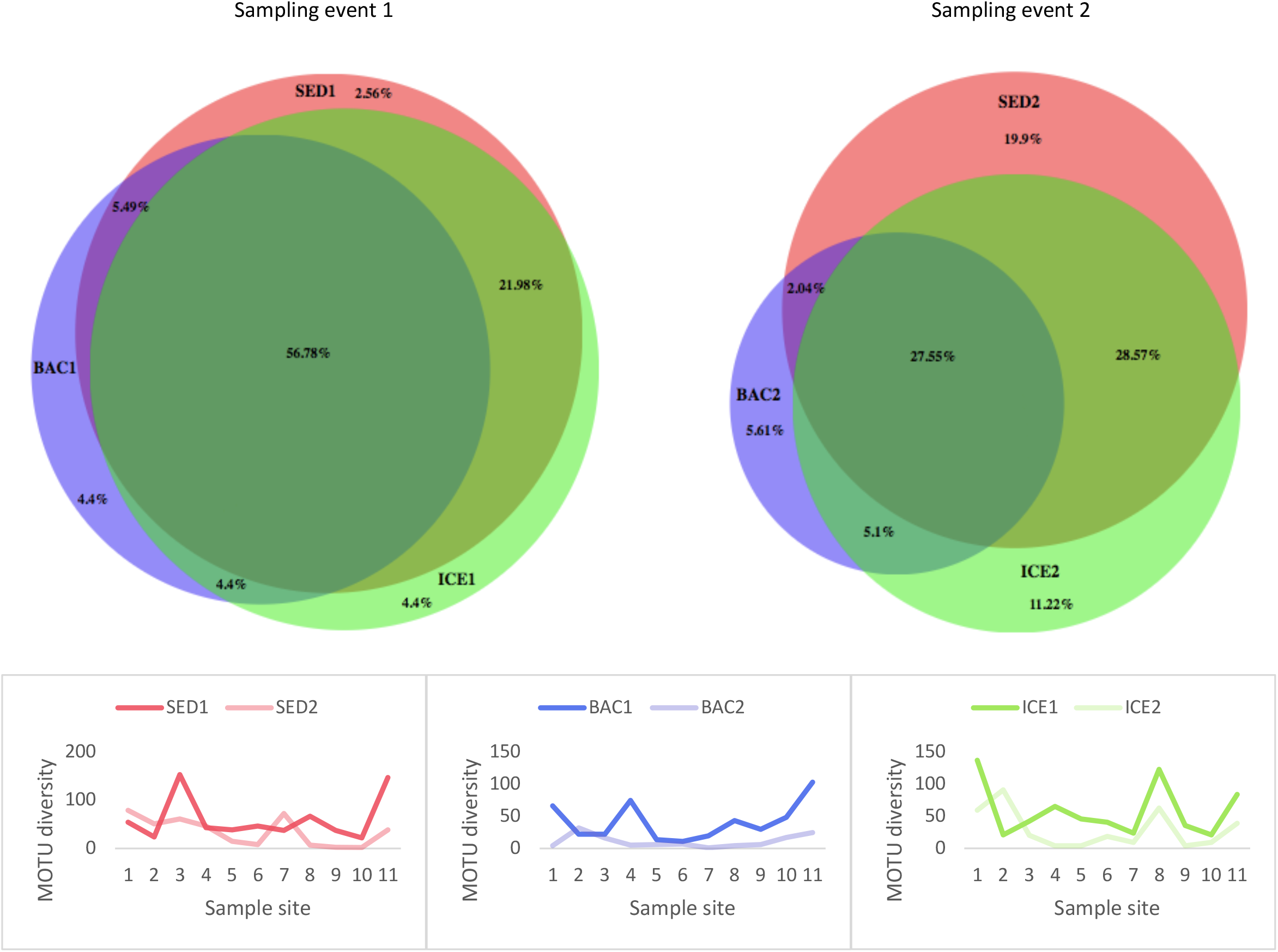
Comparison of MOTU recovery between sampling events.

### Community composition across treatments

In total, we detected 7 orders (Characiformes, Cichliformes, Clupeiformes, Cypriniformes, Cyprinodontiformes, Gymnotiformes, and Siluriformes) and 20 families. Order and family richness obtained were compared using ggplot charts (Figure 5) and showed a slight difference across all treatments. As for preservation methods, the relative read abundance (%) was similar between water samples preserved in BAC and ICE for the first sampling, however, eDNA from two families of Siluriformes (Callichthyidae and Auchenipteridae) was not recovered from samples preserved using the cationic surfactant.

**Figure 5:**
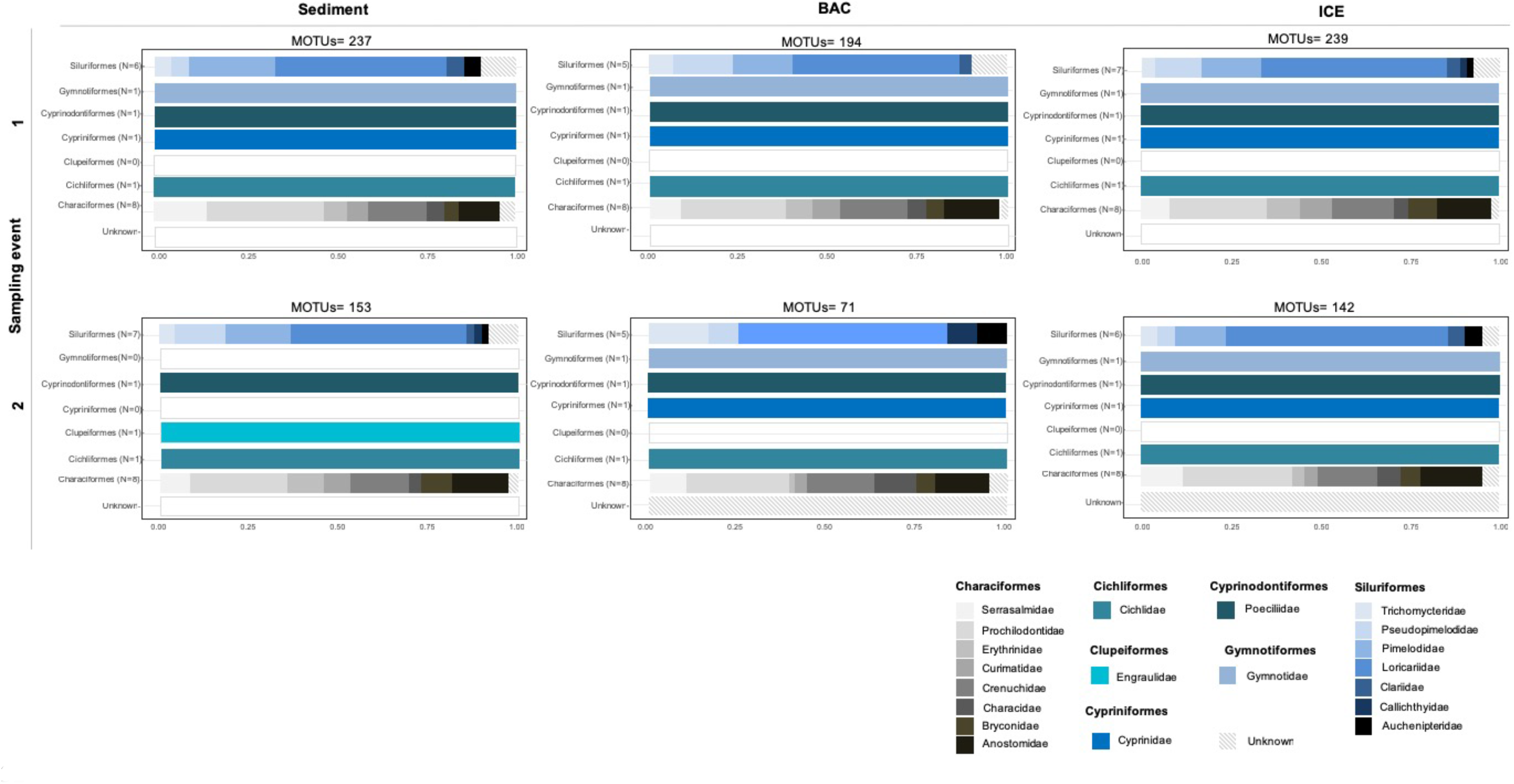
Relative read abundance per order and family.

During the second sampling, the relative read abundance slightly differed between these two methods with a highest amount of reads from Trichomycteridae (Order Siluriformes) and also absence of reads from Pimelodidae (Order Siluriformes) in samples with added BAC. Thus, samples stored in ICE outperformed samples preserved with BAC in both MOTUs recovery and order/family richness.

Regarding the sampling medium, sediment samples provided similar results to water samples, except in the order Siluformes, where it outperformed water samples preserved with BAC by detecting the family Auchenipteridae, and was surpassed by water samples preserved in ICE in detecting the family Callichthyidae, during the first sampling event. Whereas during the second sampling, the sediment samples did not recover MOTUs from two orders (Gymnotiformes and Cypriniformes) but detected one order (Clupeiformes) not identified in the water samples.

In contrast with results obtained for MOTUs recovery, despite showing a lower amount of MOTUs when compared to samples obtained in the first sampling event, samples obtained in the second event allowed the detection of additional orders and families. For the sediment samples, two orders were not detected (Cypriniformes and Gymnotiformes) but one order (Clupeiformes) and one additional family of Siluriformes (Callichthyidae) were only detected in sediments collected at the second sampling time. Regarding the samples preserved in BAC, two families of the order Siluriformes were not detected during the second sampling (Claridae and Pimelodidae) and two additional families of the same order were included (Callichthyidae and Auchenipteridae), while samples stored in ICE detected one fewer family (Callichthyidae) when compared to the first sampling.

## DISCUSSION

Despite the exponential increase of eDNA publications, most of the studies have been conducted in temperate regions and in fairly well accessible areas. To date, few studies have tested the use of eDNA metabarcoding in remote tropical sites, and to our knowledge no study encompassing freshwater fish biodiversity at a large scale has been performed in Brazil (though Cilleros et al., 2018 recently published a similar study on fish diversity of French Guiana). Here, we tested two preservation methods for water samples (cooling the samples vs adding a cationic surfactant as preservative) and also, we tested the influence of sampling medium (water vs sediment) and time on eDNA recovery to evaluate the most suitable method and provide a framework for downstream studies in tropical catchments.

Overall, comparisons between preservation methods showed a smaller effect on eDNA recovery than sampling medium and time (Table S2). Sediment and water samples kept in cooling boxes outperformed water samples preserved with the cationic surfactant solution (237 and 239 against 194 MOTUs, respectively), while the highest amount of MOTUs was detected during the first sampling event for all treatments. Most of the variance found resides within the treatments analyzed, this variance may be due to: i) the distribution of eDNA might be heterogeneous in rivers showing different spatial structures (Hänfling et al., 2016); ii) eDNA transport distances may vary between species (Deiner & Altermatt, 2014); iii) natural differences found in community composition across samples sites, as the structure of freshwater fish communities are influenced by complex interactions and by heterogeneity of freshwaters along the river gradient (e.g. geomorphic and hydrologic conditions, microbiota, temperature, pH, acidity, and chemical composition) (Spurgeon, Pegg, Parasiewicz, & Rogers, 2018). Also, as shown by Macher and Leese (2018) community composition can change even when sampling the same location in a time frame shorter than one minute and our findings also agree with earlier authors in that patterns of persistence of eDNA in rivers can be irregular.

Despite showing a significant difference, a small effect size was found for comparisons between preservation methods. The effect of preservation method might be related to the physical state of DNA molecules in the sample, free DNA can bind to humic substances and thus, be protected from enzymatic degradation and show a decreased rate on eDNA removal (Crecchio & Stotzky, 1998). Environmental DNA persistence can also be affected by the trophic state, showing a higher detectability in dystrophic and eutrophic waters than in oligotrophic systems (Eichmiller et al., 2016). The Jequitinhonha River is characterized by acid waters and contains mostly dystrophic and eutrophic soils (Intertechne, 2010) and perhaps, in this case, low temperatures could better preserve the eDNA molecules on water samples and might be more important to eDNA preservation than adding the cationic surfactant. However, degradation rates at complex tropical environments, such as the Jequitinhonha River, have not been evaluated and the trends for eDNA persistence remain unknown in this realm. A similar result was found by Laddel et al. (2018), who compared lowering the temperature of samples to adding EtOH–NaAc, where cooling of the samples outperformed the use of a buffer solution. It should also be noted that some of the discrepancies between ICE and BAC detections may simply be due to the reduction of stochasticity afforded by the additional PCRs conducted on the each water sample (six in total) (Leray & Knowlton, 2017).

Thus, despite increasing the equipment need, cooling may be considered as the first option to decrease DNA degradation in water samples during field collection. Unless no other option is available, cationic surfactant solutions might not be worthwhile for field sampling in remote areas due to the difficulties in accessing these specific laboratory reagents and the significant safety hazard posed by these chemicals (Ladell et al., 2018). However, if neither filtering nor cooling is feasible for a few hours after sampling, the use of some form of preserving buffer should remain a requirement.

Community composition is expected to differ between sampling media, as previous eDNA studies have found sediment to show a higher DNA concentration and a longer detectability than surface water (Turner et al., 2015). Since DNA can persist longer when incorporated into the sediment, temporal inference may be challenging (Turner et al., 2015); on the other hand, a higher degradation rate and lower detection lag time in aqueous eDNA samples provide a contemporary snapshot of the biodiversity being assessed (Hansen et al., 2018). Here, we have found a significant difference (p<0.05) and a higher size effect (R^2^=0.06-0.08) on MOTU recovery between sediment and water samples (Table 3). Sediment samples outperformed water samples preserved with BAC by detecting the family Auchenipteridae (Order Siluriformes), and was surpassed by water samples preserved in ICE in detecting the family Callichthyidae, during the first sampling event. In the second sampling event, sediment samples failed to detect the family Callichthyidae and the orders Gymnotiformes and Cypriniformes, however, the order Clupeiformes was only found using this type of sample, and 19.9% of the MOTUs obtained for the second sampling event was exclusive to this sampling medium. MOTUs detected only in water samples might indicate the contemporary presence of those while their absence in sediments samples may be due to a short time frame for those to settle and bind to the substrate. MOTUs belonging to the order Clupeiformes were detected only in sample site 11, located at the river mouth and refer to marine species that occasionally venture into the river to feed (Andrade-Neto, 2010). Although these species might not have been there at the time of sampling, they might have shed DNA during their incursions and the eDNA bound to sediment can have persisted longer than the eDNA in the surface water, contributing to its later detection. Thus, combining sediment and water samples may contribute to obtain a snapshot of the fish community that can distinguish between resident and transient species.

**Table 3:**
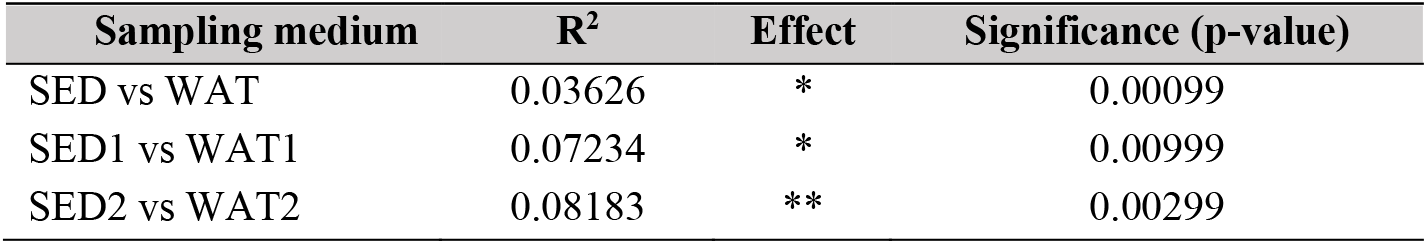
PERMANOVA results (R^2^-effect sizes and significance level) showing the effect of sampling medium on MOTU diversity recovery.

Sampling time influenced MOTU recovery and community composition in all treatments analyzed, showing a highest effect size in sediment samples and a lowest effect size in water samples preserved in BAC. An association between the number of MOTUs and effect size was found, as the higher amount of MOTUs obtained, the higher was also the effect size of sampling event. Despite showing a lower amount of MOTUs detected, samples obtained in the second event allowed the detection of additional orders and families. During the second sampling event 19.9% of the MOTUs were only detected in sediment samples when contrasted to 2.56% in the first sampling. Sediments can act as eDNA molecules reservoirs, since eDNA can settle and bind to the substrate and when incorporated its persistence can be much longer (Eichmiller et al., 2014; Turner et al., 2014).

Environmental DNA concentration can change seasonally, as well as changes in community composition over time should be expected due to natural (e.g. environmental changes, such as variation in water temperature and flow) or anthropogenic factors (e.g. pollution, introduction of physical barriers) and this variation has already been documented through metabarcoding in estuaries (Stoeckle, Soboleva, & Charlop-Powers, 2017), lakes (Bista et al., 2017) and rivers, even over a small temporal scale (Macher & Leese, 2018). The Jequitinhonha Valley is a dry region that is under the risk of desertification and by the beginning of 2017, when the first sampling event was undertaken, it was facing the worst drought in the past 80 years. However, the sampling was conducted during the rainy season and the average accumulated rainfall increased from 2.1-50mm (first sampling time) to 100-250 mm (second sampling event) per month (CPTEC/INPE, 2018). The increase in the precipitation level in this region, with heavy rainfall causing floods in several sites and this seasonal change might have impacted the MOTU recovery during the second sampling, as the increase in water level can contribute to dilute the eDNA, change the water temperature and flow, and also cause fluctuations in community composition. Increased water volume after the rainfall contributes to a higher velocity and affects eDNA concentrations in water columns, as eDNA molecules are transported and dispersed towards downstream river (Shogren et al., 2017). Furthermore, an increase in water flow caused by rainfall might lead to eDNA particles resuspension, which could explain a higher similarity detected by the nMDS between sampling medium in the second sampling event.

Understanding the effect of abiotic and biotic factors on eDNA recovery in tropical lotic environments is crucial to improve the interpretation of results and assure the effectiveness of eDNA as a biodiversity assessment tool. Here, we showed the first results on effect of sampling medium, time, and preservation methods in lotic environments and our findings suggest that the interaction between preservation method and MOTU recovery might be less significant than the influence of sampling medium and sampling event. Cooling the water samples before the filtering might be a better option in field work conducted in remote areas due to logistical issues and to an increased eDNA recovery when compared to addition of cationic surfactants as sample preservatives.

We also highlight the importance of a better interpretation of eDNA results when comparing sediment and water samples due to distinct temporal intervals covered, and comparing two sets of samples obtained in a short time interval we demonstrate the importance of applying multiple sampling collections when planning a realistic screening of fish biodiversity in lotic environments. The recovery of a high amount of MOTUs allowed the detection of a high degree of fish biodiversity, including changes in community composition, demonstrating the effectiveness of eDNA as a biodiversity assessment tool in neotropical lotic rivers. However, this study was method-focused and detailed ecological analysis of the recovered biodiversity is the natural next step. This will require an improved reference database, as the data obtained here (i.e. potentially hundreds of fish species) suggests that the biodiversity of this catchment is grossly underestimated (Andrade-Neto, 2010).

## Supporting information

## ACKNOWLEDGEMENTS

This work was supported by Conselho Nacional de Desenvolvimento Científico e Tecnológico (CNPq) and Science without Borders Program (Grant #204620/2014-7). We are grateful to Letícia Sales for the invaluable fieldwork assistance, Charles Baillie for the advice on bioinformatics analysis, and Gilberto Salvador for his assistance with the maps.

## DATA ACCESSIBILITY

Data will be made public on the DRYAD repository upon acceptance.

## AUTHOR CONTRIBUTIONS

Study design: NGS and SM. Field work and sample collection: NGS. Laboratory experiment: NGS and OSW. Data analyses: NGS, OSW, SM. Manuscript writing: NGS, DCC, OSW, and SM.

