## Supplementary material for "Influence of Preservation Methods, Sample Medium and Sampling Time on eDNA Recovery in a Neotropical River"

**APPENDICES AND SUPPLEMETARY MATERIALS**

Table S1: Sample sites, including GPS coordinates.

| ID | CODE | Sample site | GPS Coordinates | |
|---|---|---|---|---|
| 1 | MED | Medanha | 18° 7'15.06"S | 43°30'59.16"W |
| 2 | TB | Terra Branca | 17°18'48.34"S | 43°12'26.61"W |
| 3 | JGON | José Gonçalves (upstream the UHE Irapé dam) | 16°44'25.89"S | 42°34'16.34"W |
| 4 | ITAC | Itacambiruçu | 16°36'24.00"S | 42°49'46.00"W |
| 5 | CM | Coronel Murta (downstream the UHE Irapé dam) | 16°44'26.85"S | 42°34'11.78"W |
| 6 | ARA | Araçuaí | 16°51'10.47"S | 41°51'33.53"W |
| 7 | JEQ | Itaobim/Jequitinhonha | 16°26'16.74"S | 41°1'1.45"W |
| 8 | ALM | Almenara/Jacinto | 16° 8'26.20"S | 40°35'4.64"W |
| 9 | SD | Salto da Divisa | 15°59'51.07"S | 39°53'29.76"W |
| 10 | ITAP | Itapebi | 15°56'57.69"S | 39°31'27.08"W |
| 11 | BEL | Belmonte | 15°51'0.02"S | 38°52'13.66"W |

Table S2: PERMANOVA results ($R^2$-effect sizes and significance level) showing the effect of preservation method, sampling medium, and sampling time on MOTU diversity recovery

| | $R^2$ | Effect | Significance (p-value) |
|---|---|---|---|
| **Preservation method** | | | |
| BAC1 vs ICE1 | 0.02636 | * | 0.01698 |
| BAC2 vs ICE2 | 0.02780 | * | 0.06493 |
| **Sampling medium** | | | |
| SED vs WAT | 0.03626 | * | 0.00099 |
| SED1 vs WAT1 | 0.07234 | * | 0.00999 |
| SED2 vs WAT2 | 0.08183 | ** | 0.00299 |
| SED1 vs BAC1 | 0.06006 | * | 0.00099 |
| SED1 vs ICE1 | 0.05980 | * | 0.00099 |
| SED2 vs BAC2 | 0.08410 | ** | 0.00099 |
| SED2 vs ICE2 | 0.07056 | * | 0.00099 |
| **Sampling time** | | | |
| SED1 vs SED2 | 0.07762 | * | 0.00099 |
| BAC1 vs BAC2 | 0.04192 | * | 0.00099 |
| ICE1 vs ICE2 | 0.06436 | * | 0.00099 |
